# The mid-lateral cerebellum is necessary for reinforcement learning

**DOI:** 10.1101/2020.03.20.000190

**Authors:** Naveen Sendhilnathan, Michael E. Goldberg

**Affiliations:** Doctoral program in Neurobiology and Behavior, Columbia University, New York, NY; Dept. of Neuroscience, Mahoney Center for Brain and Behavior Research, Zuckerman Mind, Brain, and Behavior Institute, Columbia University, New York, NY; New York State Psychiatric Institute, New York, NY; Kavli Institute for Brain Science, Columbia University, New York, NY; Dept. of Neurology, Psychiatry, and Ophthalmology, Columbia University College of Physicians and Surgeons, New York, NY

## Abstract

The cerebellum has long been considered crucial for supervised motor learning and its optimization^1-3^. However, new evidence has also implicated the cerebellum in reward based learning^4-8^, executive function^9-12^, and frontal-like clinical deficits^13^. We recently showed that the simple spikes of Purkinje cells (P-cells) in the mid-lateral cerebellar hemisphere (Crus I and II) encode a reinforcement error signal when monkeys learn to associate arbitrary symbols with hand movements^4^. However, it is unclear if the cerebellum is necessary for any process beyond motor learning. To investigate if the mid-lateral cerebellum is actually necessary for learning visuomotor associations, we reversibly inactivated the mid-lateral cerebellum of two primates with muscimol while they learned to associate arbitrary symbols with hand movements. Here we show that cerebellar inactivation impaired the monkey’s ability to learn new associations, although it had no effect on the monkeys’ performance on a task with overtrained symbols. A computational model corroborates our results. Cerebellar inactivation increased the reaction time, but there were no deficits in any motor kinematics such as the hand movement, licking or eye movement. There was no loss of function when we inactivated a more anterior region of the cerebellum that is implicated in motor control. We suggest that the mid-lateral cerebellum, which provides a reinforcement learning error signal^4^, is necessary for visuomotor association learning. Our results have implications for the involvement of cerebellum in cognitive control, and add critical constraints to brain models of non-motor learning^14,15^.

---

We trained two monkeys to associate arbitrary visual symbols with left and right hand movements to earn an immediate liquid reward. On each recording session, the monkeys first performed 30-50 trials of the task with familiar, overtrained symbols (**Fig 1a**) at a success rate close to 100%. We created a library of 16 pairs of fractal symbols with varying levels of similarities between the symbols. After the monkeys performed a few trials on the overtrained task (OT), we presented them with one of the novel symbol pairs from the library (**Fig 1a**) and tracked their learning. To calculate the monkeys’ acquisition-difficulty level for different symbol pairs, we recorded the monkeys’ performance for each pair (Monkey B-16 pairs, Monkey S-9 pairs) at least three times, spread over several weeks, presented in random order (**Fig 1b**). To prevent the monkeys from relying on previous experience with the same symbol pair to relearn their associations, we either presented a given symbol pair again only several days after its prior presentation, with intervening presentations of other symbol pairs or random pairs not from the library (**Fig 1b;** symbol pair highlighted in yellow), or we reversed the symbol-hand association the next time we presented the same pair (**Fig 1b;** symbol pair highlighted in red). We collected three repetitions of behavioral performance for each symbol pair from both monkeys (Monkey B-16 pairs, Monkey S-9 pairs). We defined the acquisition-difficulty level for a given symbol pair, for a specific monkey, as the trial at which the monkeys’ performance was at least 75% accurate (see methods). The acquisition-difficulty rates were similar for each presentation of a given symbol pair (**Fig 1c**) indicating that the monkeys inevitably forgot their prior experience with a given symbol pair during each additional re-learning.

**Figure 1:**
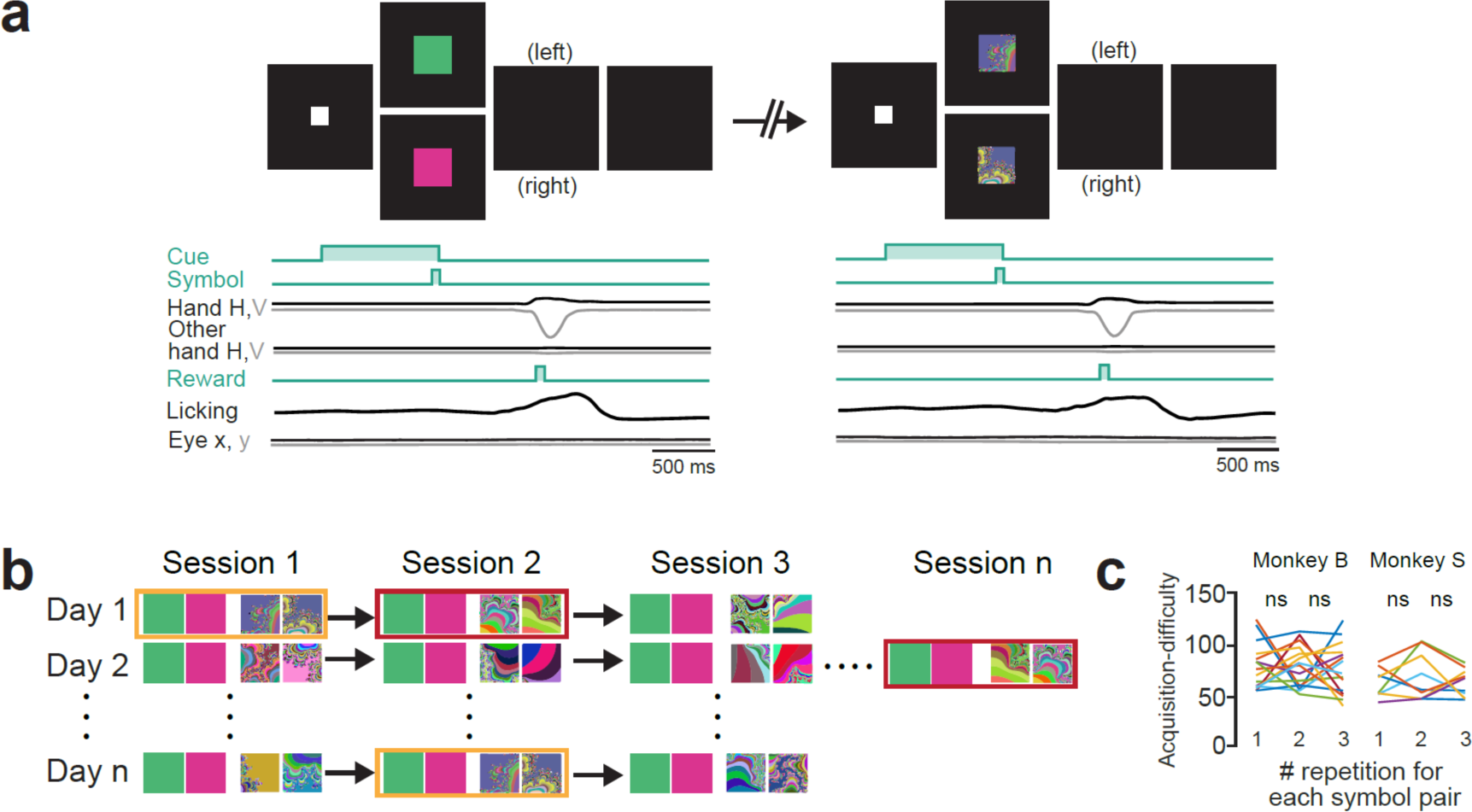
Visuomotor association learning task paradigm. **a.** Two-alternative forced-choice discrimination task (top) and parameters (bottom): The monkeys pressed each bar with a hand, and a white square appeared that served as the cue for the start of the trial. Then one of the two visual symbols appeared briefly. The monkey lifted the hand associated with the presented symbol to earn a reward immediate. Left: a trial from overtrained condition and right: a trial from novel learning condition. **b.** Trial session organization. Sessions marked with yellow rectangle indicate repeated presentation of the same symbol pairs. Sessions marked with red rectangle indicate repeated presentation of the same symbol pairs but with reversed visuomotor association. **c.** Acquisition-difficulty for each symbol pair (colored lines) per repetition, for both monkeys. Acquisition-difficulty for the second learning of the same symbol pair was not different from the first learning for either monkeys (monkey B: P = 0.8608; monkey S: P = 0.5489 paired t-test). Similarly, Acquisition-difficulty for the third learning was not different from the second learning for either monkeys (monkey B: P = 0.2640; monkey S: P = 0.5228 paired t-test).

Although P-cells in the mid-lateral cerebellum, encode a reinforcement learning error signal^4^, it is not clear if this activity is only correlative or causal to behavior. To test if mid-lateral cerebellum is necessary for the monkeys to perform the task, we infused 4-10 μl of a 10 mg/ml solution of muscimol, a GABA_A_ agonist that hyperpolarizes cell bodies without affecting fibers of passage, near crus I and II of the mid-lateral cerebellum of two monkeys in the same location where we had recorded task-dependent P-cells (see methods, **Fig 2a;** see **Fig S1** for infusion locations). We infused saline alone on the days before and after every muscimol injection as a control.

**Figure 2:**
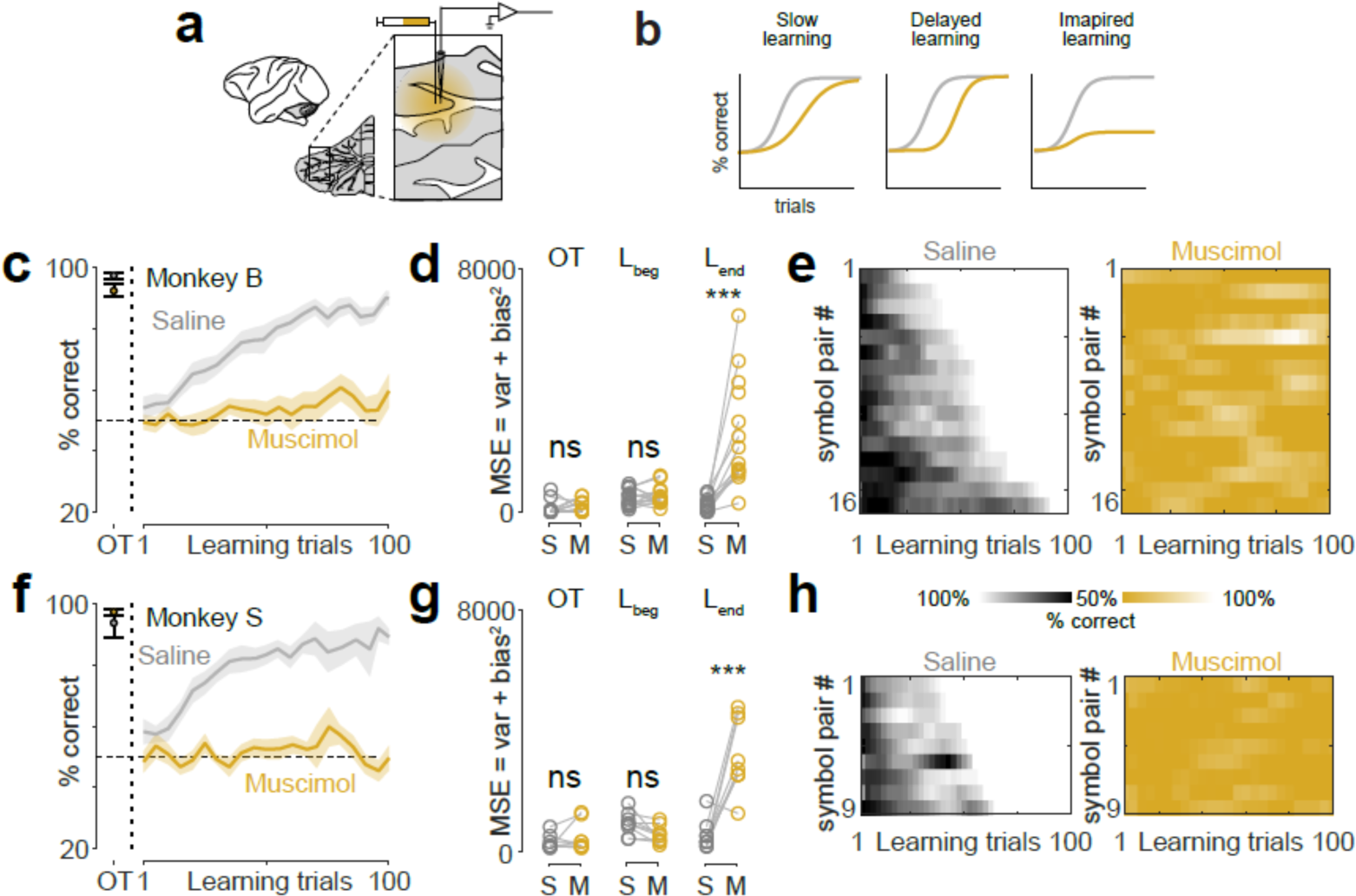
Cerebellar inactivation caused an impairment in visuomotor association learning. a. Schematic illustatrion of the mid-lateral cerebellum inactivation and simltaneous recording. b. Schematic illustation of three possible ways in which cerebellar inactiavtion (yellow) can affect the learning curve compared to control conditions (gray). c. Behavioral performace of Monkey B during OT and learning trials for control-saline (gray) and cerebellar inactivation (yellow) conditions. d. Quantitation of mean squared error (MSE = bias^2^ + variance) from **Fig 2c**; OT: P = 0. 0.2518, ranksum test; L_beg:_ P = 0. 1117, paired t-test; L_end_: P = 2.6410e-05.,paired t-test. For the bias^2^ and variance, see **Fig S2**. e. Left: heat plot of learning curves (each row) for the behavioral performaces of 16 pairs of symbols, arranged in the increasing oder of acquisation difficulty during the saline-control condition. Right: heat plot of learning curves (each row) for the behavioral performaces of the same16 pairs of symbols (in the same oder of left panel) during cerebellar inactivation. f. Same as **Fig 2c**, but for monkey S. g. Qauntitation mean squared error (MSE) from **Fig 2g**; OT: P = 0.2787, ranksum test; L_beg:_ P = 0.2196, paired t-test; L_end_: P = 0.0016, ranksum test. For the bias^2^ and variance, see **Fig S2**. h. Same as **Fig 2e**; but for Monkey S.

The monkeys typically learned the novel association in 50-70 trials during the control sessions (**Fig 2c, 2f**). Mid-lateral cerebellar inactivation prevented them from learning the correct association for the same pairs of symbols from the library that we used for the saline condition. In the beginning of learning, the behavioral performances for control and inactivation conditions were comparable, with low mean squared error (Monkey B: P = 0.1117; t-test; **Fig 2d;** Monkey S: P = 0.2196; paired t-test; **Fig 2g**). However, after 75 trials of learning, when the monkeys performed more than 75% correct with low mean squared error during control conditions, their performance was close to chance level (Monkey B: P = 2.6410e-05; t-test; **Fig 2d;** Monkey S: P = 4.1135e-05; ranksum test; **Fig 2g**) with significantly higher mean squared error during the inactivation condition (Monkey B: P = 3.3216e-04; ranksum test; **Fig 2d;** Monkey S: P = 0.0016; ranksum test; **Fig 2g**). Cerebellar inactivation affected the bias more than the variance during learning (**Fig S2**). The learning performance during cerebellar inactivation was independent of the rate of learning of a symbol pair in the control condition for both monkeys. When the cerebellum was inactivated, the monkeys could not even learn the association that had the least acquisition difficulty (**Fig 2e, 2h**). Repetition of symbols after error trials during learning significantly improved the performance (rate of learning) under control-saline condition (P = 0.0029; paired t-test) but not during cerebellar inactivation (P = 0.1117; paired t-test) for the same number of trials, for the same symbol pairs (**Fig S3**). We verified that the performance modulations across sessions, within a day, did not confound our observation (**Fig S4**).

Mid-lateral cerebellar inactivation did not alter the monkeys’ behavioral performance on the overtrained task: The monkeys continued to perform close to 100% accuracy when the cerebellum was inactivated, similar to their performance during the saline-control condition (Monkey B: P = 0.2110; Mann-Whitney U test; **Fig 2c;** Monkey S: P = 0.9132; Mann-Whitney U test; **Fig 2f**). The inactivation did not affect the mean squared error (Monkey B: P = 0.2518; Mann-Whitney U test; **Fig 2d;** Monkey S: P = 0.2787; Mann-Whitney U test; **Fig 2g**) in the performance, the decomposed bias squared (**Fig S2**) or the variance (**Fig S2**) during the overtrained task.

Although inactivation of mid-lateral cerebellum prevented the monkeys from learning the visuomotor association for a given hand movement, it did not affect the gross motor kinematics of hand movement: In both control and inactivation conditions, the monkeys performed the task with well-stereotyped hand movements similar to those in the OT (**Fig 3a;** Monkey B: P = 0.7342; Mann-Whitney U test; Monkey S: P = 0.5286; paired t-test) and during learning (**Fig 3b;** Monkey B: P = 0.8703; paired t-test; Monkey S: P = 0.3195; paired t-test). The licking behavior was also not different between the control and inactivation conditions for OT (**Fig 3c;** Monkey B: P = 0.7631; paired t-test; Monkey S: P = 0.9965; paired t-test) or during learning (**Fig 3d;** Monkey B: P = 0.4413; paired t-test; Monkey S: P = 0.4217; paired t-test). Although monkeys generally tended to explore their environment more during learning than during OT, there were no systematic differences in their eye movements between the control and inactivation conditions during OT (**Fig 3e;** Monkey B: P = 0.2280; Mann-Whitney U test; Monkey S: P = 0.7311; paired t-test) or during learning **Fig 3f;** Monkey B: P = 0.1127; Mann-Whitney U test; Monkey S: P = 0.6665; Mann-Whitney U test). Additionally, cerebellar inactivation did not have a differential effect on motor kinematics for correct and wrong trials (**Fig S6**). Furthermore, with cerebellar inactivation, the monkeys had no deficit performing the OT task with very different hand movements. We switched the-manipulanda that the monkeys used to reporting their choices, from bars to dowels. Although monkeys had to make an entirely different hand movement to report the same choices (**Fig S7**) cerebellar inactivation had no effect on the kinematics of the changed movement (**Fig S7**). The only motor effect of the inactivation was to increase monkeys’ manual reaction time for both monkeys during learning (P = 0.0013; paired t-test; **Fig 3g, h**). Given that there were no changes in any other aspect of the movements with which the monkeys signaled their decision, we inferred that inactivating the midlateral cerebellum interfered with the monkeys’ ability make a decision about the association than their ability to report it.

**Figure 3:**
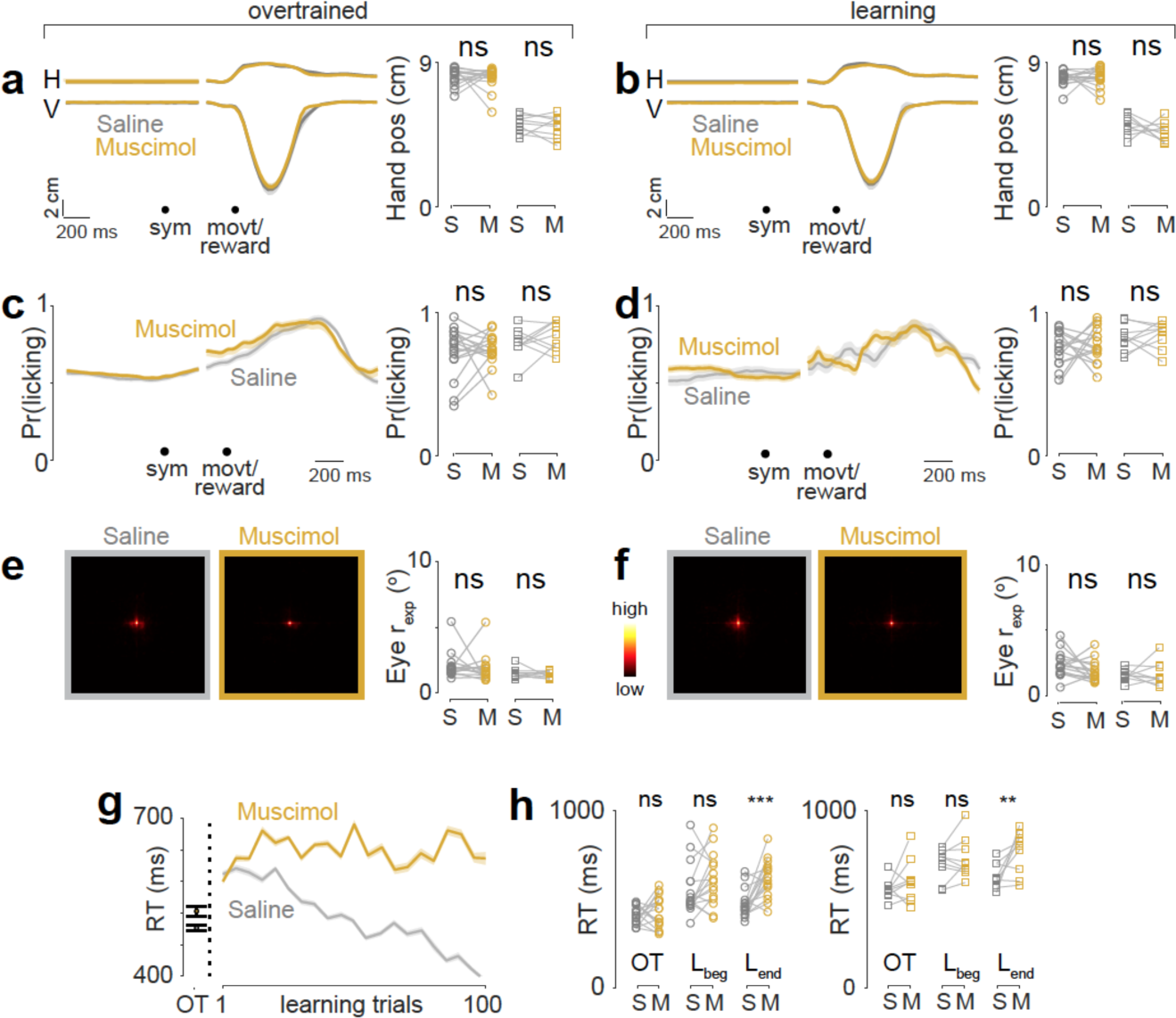
Cerebellar inactivation did not cause any deficit in motor kinematics. a. Left: Average horizontal (H) and vertical (V) hand positions across sessions for monkey B, during OT for control-saline (gray) and muscimol-inactivation (yellow) conditions. See **Fig S5** for Monkey S. Right: Quantitation of the amplitude of hand movement for monkey B (left; circle markers; P = 0.7342, ransksum test) and monkey S (right, square markers; P = 0.5286; paired t-test). b. Same as **Fig 3a**; but during learning condition. Right panel, monkey B (left; circle markers; P = 0.8703, paired t-test) and monkey S (right, square markers; P = 0.3195; paired t-test). c. Left: Average licking activity across sessions and monkeys during OT for control-saline (gray) and muscimol-inactivation (yellow) conditions. Right: Quantitation for monkey B (left; circle markers; P = 0.7631, paired t-test) and monkey S (right, square markers; P = 0.9965; paired t-test). d. Same as **Fig 3c**; but during learning condition. Right panel, monkey B (left; circle markers; P = 0.4413, paired t-test) and monkey S (right, square markers; P = 0.4217; paired t-test). e. Left: Average eye movement exploration activity on the full screen, across sessions and monkeys during OT for control-saline (gray) and muscimol-inactivation (yellow) conditions. Heat map is shown in the inset. Right: Quantitation of radius of visual exploration (r_exp_; see methods) for monkey B (left; circle markers; P = 0.2280, ranksum test) and monkey S (right, square markers; P = 0.7311; paired t-test). f. Same as **Fig 3e**; but during learning condition. Right panel, monkey B (left; circle markers; P = 0.1127, ranksum test) and monkey S (right, square markers; P = 0.6665; ranksum test). g. Average RT profile during OT and learning across sessions and monkeys for control-saline (gray) and inactivation (yellow) conditions. h. Quantitation from **Fig 3g** for two monkeys separately. Left: Monkey B, OT: P = 0.4774; paired t-test; L_beg_: P = 0.1011; paired t-test; L_end_: P = 3.1909e-04 paired t-test. Right: Monkey S, OT: P = 0.3028; paired t-test; L_beg_: P = 0.2826; paired t-test; L_end_: P = 0.0021 paired t-test.

Finally, to test if the effect of muscimol inactivation were limited to the mid-lateral cerebellum, we made muscimol injections into more anterior parts of the cerebellum (lobule V, an area thought to be involved in motor control^16^) in both monkeys (**Fig S9a-b**). We again recorded the monkeys’ performance for the some of the same symbol pairs from the library, used for prior experiments (Monkey B-5 pairs, Monkey S-4 pairs). Inactivation of the anterior cerebellum had no effect on the monkey’s ability to learn new visuomotor associations or to perform the overtrained association (**Fig S9c-g**). However, it caused transient hind limb ataxia and nystagmus in one monkey and slightly reduced the monkeys’ reaction time (**Fig S9e-g**).

The mid-lateral cerebellum, but not more anterior regions, lies within a closed loop network for non-motor learning, due to its anatomical connections with the basal ganglia and the prefrontal cortex^14,17-19^. Neurons in the prefrontal cortex^20-22^ the globus pallidus internal segment (GPi)^23^, caudate nucleus report the progress of visuomotor association^21^. Lesions in prefrontal cortex of the monkey eliminated the ability of monkeys to learn a new visuomotor association and impaired their ability to retain associations learned preoperatively^24^. Combined lesions of basal ganglia and premotor cortex prevented monkeys from retaining associations learned preoperatively, but did not affect their ability to learn new associations^25^.

We recently showed that the simple spikes of P-cells in the mid-lateral cerebellum encode reinforcement error signals during visuomotor association learning^4^. Two types of P-cells report whether the monkey’s decision on the most recent trial was correct (correct-preferring neurons, cP-cells) or wrong (wrong-preferring neurons, wP-cells), and although each neuron reports this only during a brief epoch (the ‘delta epoch’), the population as a whole maintains the information from shortly after the previous outcome to shortly after the next. The magnitude of this error signal is greatest at the beginning of learning, and approaches zero as the monkey learns the task, which is what is expected of a reinforcement learning error signal. The P-cells have no information about what the monkey will do in the upcoming trial or about past trials other than the most recent. When the monkeys perform the overtrained task, the neurons do not signal the occasional wrong decision. In keeping with this, reversible inactivation of this area eliminates the ability of the monkey to learn new associations, but does not affect their ability to remember the overtrained association. Inactivation increases the reaction time of the hand movement, but the absence of any other kinematic effect suggests that this increase in reaction time arises from a difficulty the monkeys have in making a decision, rather than a motor deficit per se.

A similar dissociation between short-term and chronic learning has been demonstrated in the motor system. Cats exposed to 60 minutes of exposure to .25x miniaturizing spectacles reduce the gain of their vestibuloocular reflex(VOR) to 0.75 - 0.8 from 1^26^. Injection of CQNX, an antagonist of the glutamate AMPA and kainite receptors into the flocculus, the vestibulocerebellum, eliminated this gain reduction. If the cats wore the spectacles for 3 days, and had three hour-long training sessions, their gain fell to 0.65, but CQNX injection increased it only slightly, to 0.75. These data implied that the long-term gain reduction was not exclusively dependent upon the cerebellum, but migrated elsewhere as learning proceeded. In the case of the VOR, this extracerebellar locus is the medial vestibular nucleus^27^. In the case of reinforcement learning, the synaptic changes associated with learning that begin in the cerebellum might migrate to the prefrontal cortex, the basal ganglia, or both^18^. This still needs to be investigated.

## Acknowledgments

We thank John Caban for making injectrodes and all other creative machining, Glen Duncan for highly creative and wonderful electronic assistance, Dr. Girma Asfaw, Dr. Moshe Shalev for animal care, Whitney Thomas and Holly Cline for facilitating everything. This work was supported by the Keck, Zegar Family, and Dana Foundations and the National Eye Institute (R24 EY-015634, R21 EY-017938, R21 EY-020631, R01 EY-017039, and P30 EY-019007 to M. E. Goldberg, PI).

**Raw data and computer codes are available upon request.**

---

**Figure S1:**
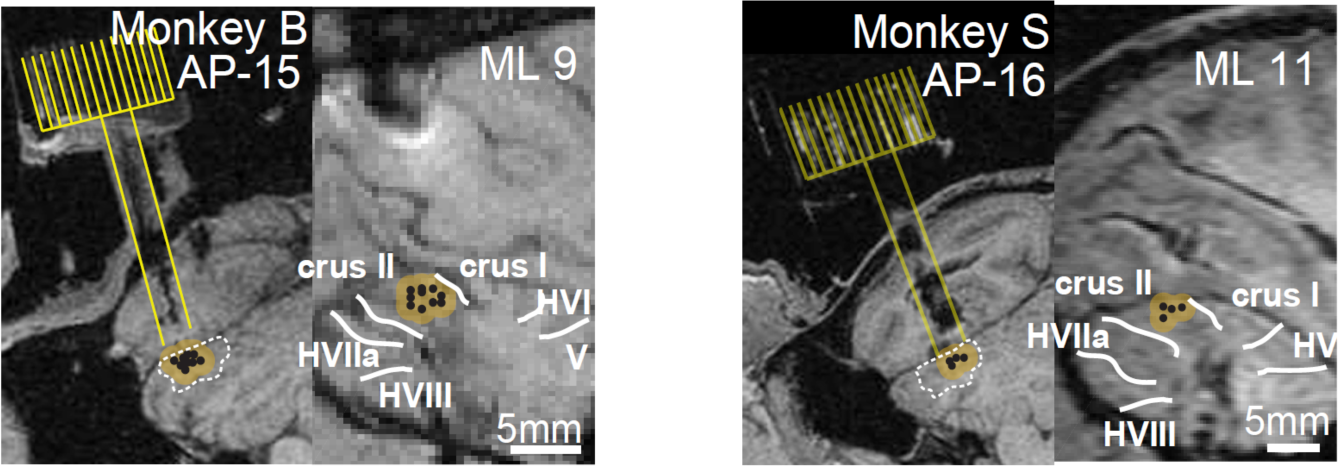
Recording and infusion locations. T1 MRI of the cerebellum infusion locations (black markers) in the coronal view (left) and sagittal view (right) for monkeys B and S. Yellow shading region surrounding the black markers represent the extent of Muscimol diffusion-spread per injection. The region marked by the white broken line is the projected region to contain P-cells that carry reinforcement error signal based on prior recordings.

**Figure S2:**
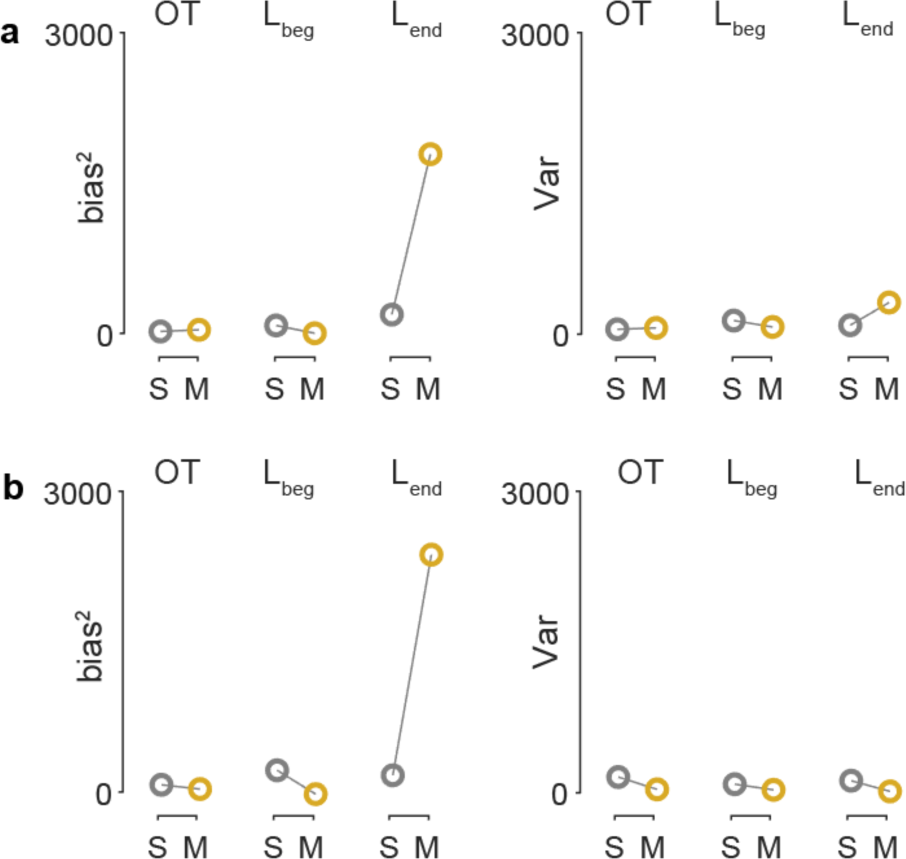
Cerebellar inactivation increased bias and variance in performance during learning but during OT condition. a. Changes in bias^2^ (left) and changes in variance (right) for Monkey B from control-saline (S, gray) to Muscimol (M, yellow) condition during OT, beginning of learning (L_beg_) and end of learning (L_end_). b. Same as **Fig S2a**, but for Monkey S.

**Figure S3:**
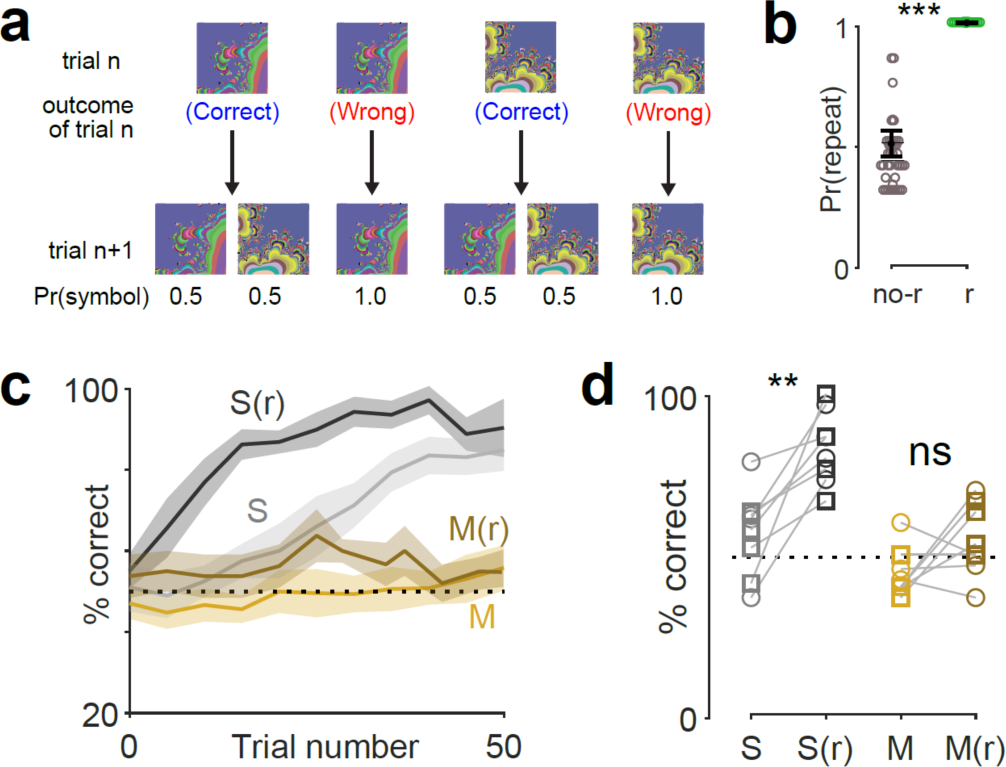
Repetition of symbols after error trials during learning significantly improved the performance during control-saline condition but not during cerebellar inactivation. a. Task paradigm: If the monkey got a trial wrong, the same symbol was presented again in the next trial (repeat condition); else, one of the two symbols was presented with equal probability in the next trial. b. Beeswarm plot of the probability of repetition of the same symbol after a wrong trial, in the normal task (left) and the repeat task (right). P = 1.9603e-08; ranksum test. c. Average learning curve (from both monkeys) for saline-control (S), saline-control with repeat (S(r)), Muscimol (M) and Muscimol with repeat (M(r)) conditions. d. Quantitation from **Fig S3c** in the trial window of 10-20 trials, for individual sessions, for two monkeys separately (circles are Monkey B; squares are Monkey S). Repetition of symbols during learning significantly improved the performance (rate of learning) during control-saline condition (P = 0.0029; paired t-test) but not during cerebellar inactivation (P = 0.1117; paired t-test).

**Figure S4:**
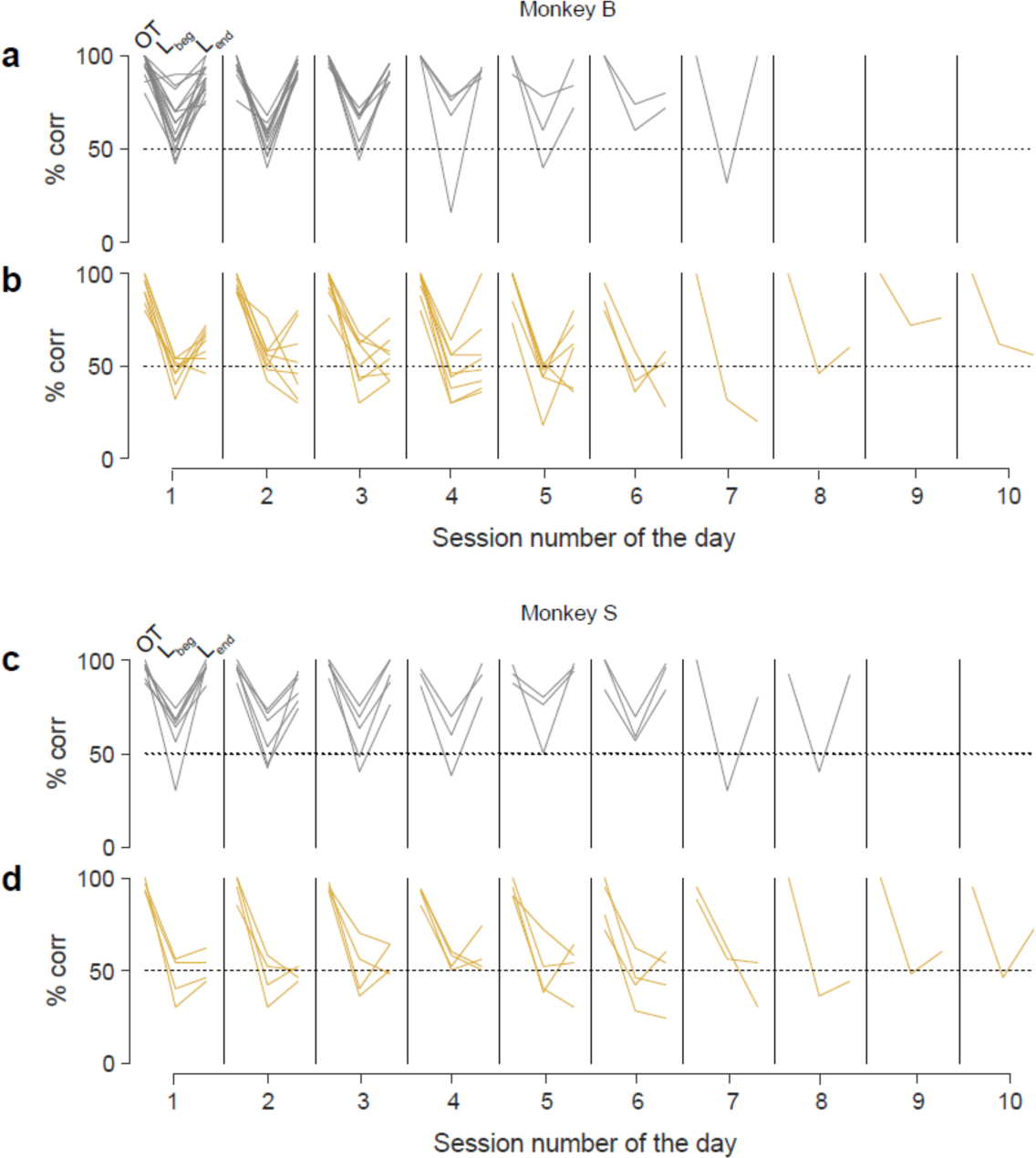
Performance within a day for different learning sessions. a. Behavioral performance for monkey B during saline-control condition across sessions for each day, for OT, beginning of learning (L_beg_) and end of learning (L_end_). b. Same as **Fig S4a**, but during cerebellar inactivation condition. c. Same as **Fig S4a**, but for monkey S. d. Same as **Fig S4b**, but for monkey S.

**Figure S5:**
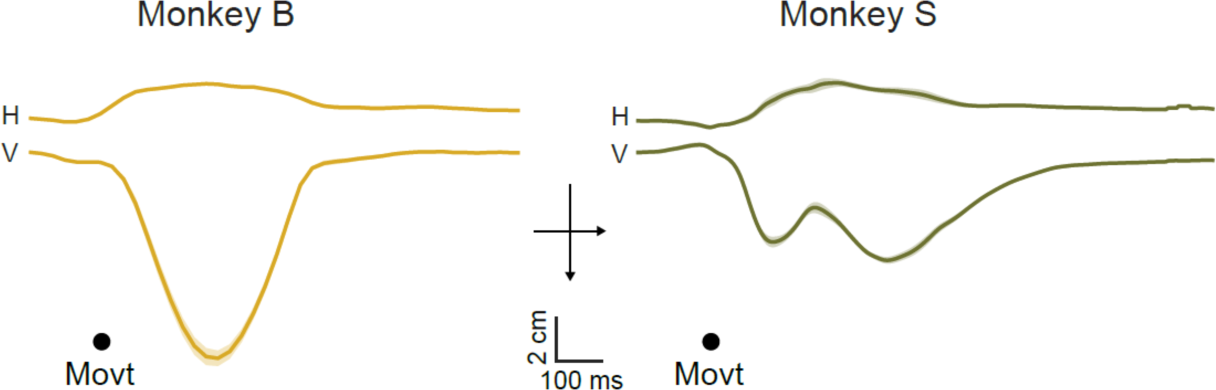
Hand positions for both monkeys for bar release condition. Horizontal (H) and vertical (V) hand positions triggered on the movement onset, for Monkey B (left) and Monkey S (right) during OT task for bar release movement.

**Figure S6:**
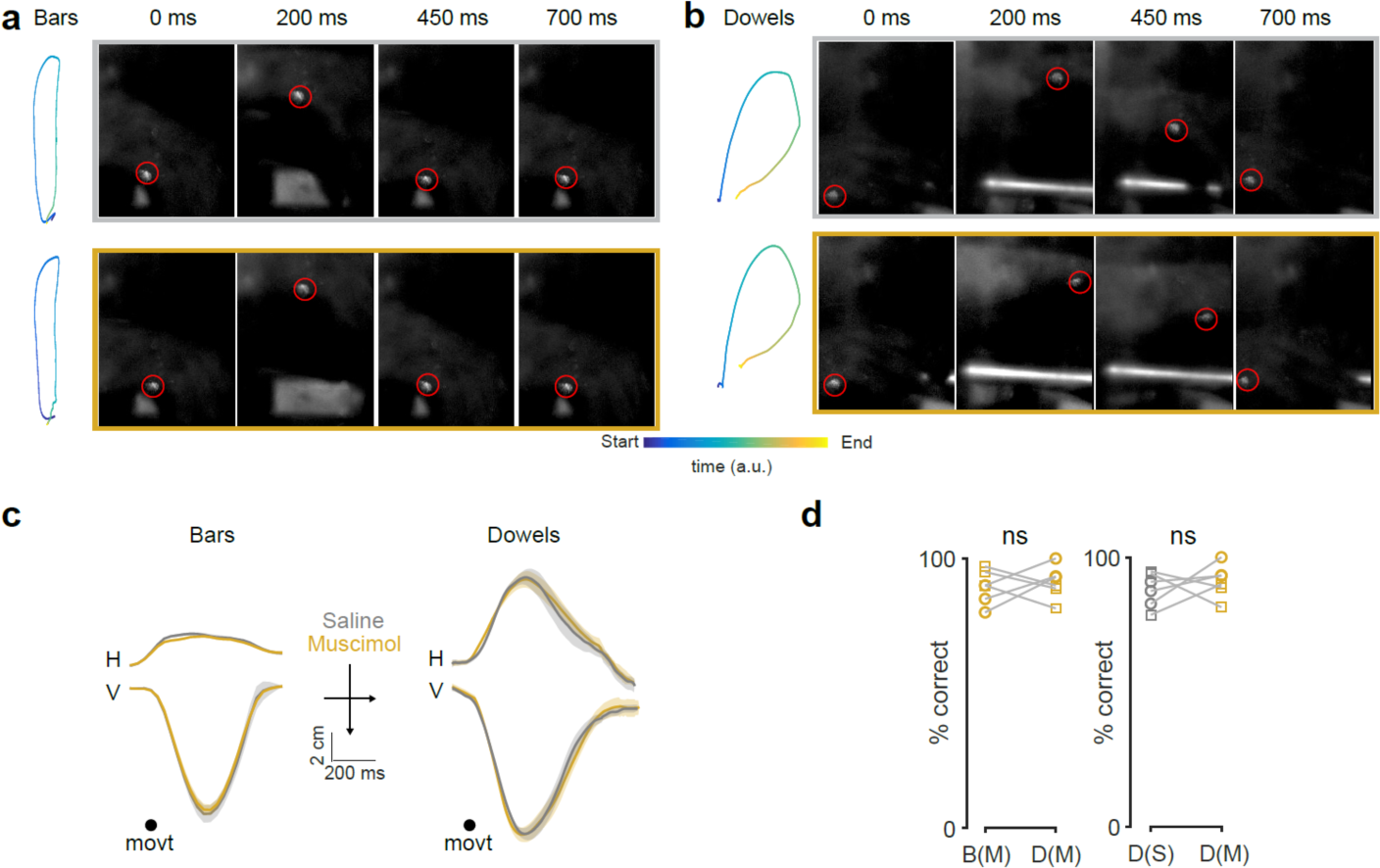
Cerebellar inactivation did not impair the ability to make different hand movements. a. Top left: Hand position in space and time (color indicates the time as in the color bar inset) for a representative trial during saline condition. Right: Snapshots from high-frame rate movies showing the monkey’s hand movement trajectory at four time points (0, 200 and 450 and 700 ms from the start of movement) for bar release. Red circle highlights the fluorescent marker. Bottom: same as top, but for a representative trial during Muscimol condition. b. Same as a, but for dowel release condition. c. Left: Average horizontal (H) and vertical (V) hand trajectories for bar release, aligned on movement onset during saline (grey) and Muscimol (yellow) conditions. Right: Same as left but for dowels condition. d. Behavioral performance of monkeys (monkey B: circles, monkey S: squares) during overtrained condition for bars vs dowels under Muscimol (left; P = 0.3744; paired t-test); and between dowels for saline and Muscimol (right; P = 0.7009; paired t-test).

**Figure S7:**
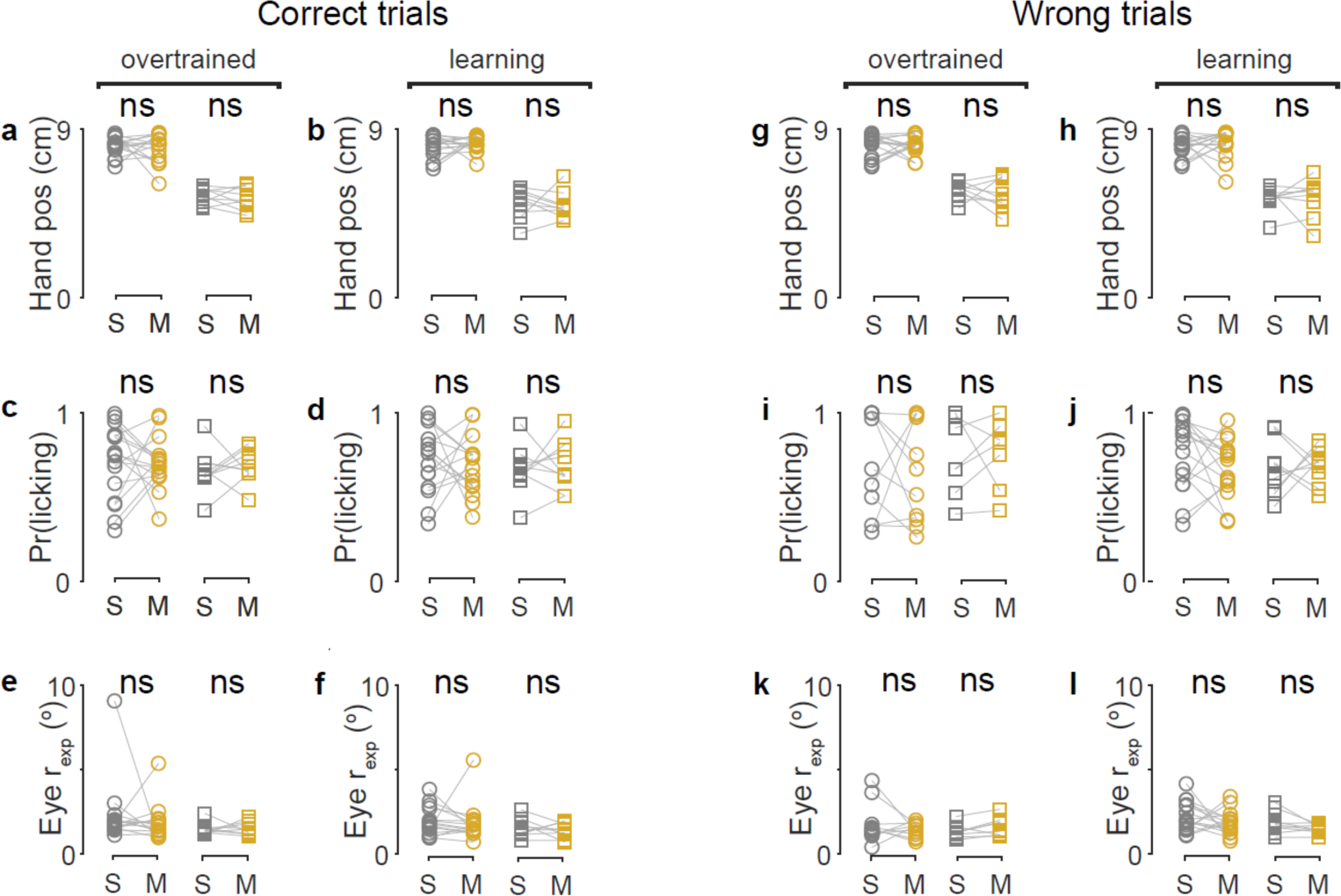
Cerebellar inactivation did not have differential effects on motor parameters for correct and wrong trials during learning. a. Amplitude of hand position on correct trials, OT, Monkey B (left): P = 0.9849; ranksum test; monkey S (right), P = 0.5362; paired t-test b. Amplitude of hand position on correct trials, learning, Monkey B (left): P = 0.4614; paired t-test; Monkey S (right): P = 0.5307; ranksum test c. Probability of licking on correct trials, OT, Monkey B (left): P = 0.9176; paired t-test; Monkey S (right): P = 0.6318; paired t-test d. Probability of licking on correct trials, learning, Monkey B (left): P = 0.6636; t-test; Monkey S (right): P = 0.7334; paired t-test e. Visual exploration on correct trials, OT, Monkey B: P = 0.9527; ranksum test; Monkey S: P = 0.9999; ranksum test f. Visual exploration on correct trials, learning, Monkey B: P = 0.9098; ranksum test; Monkey S: P = 0.4624; paired t-test g. Hand position on wrong trials, OT, Monkey B (left): P = 0.6509; ranksum test; Monkey S (right): P = 0.5833; paired t-test h. Hand position on wrong trials, learning, Monkey B (left): P = 0.1989; ranskum test; Monkey S (right): P = 0.5877; ranksum test. i. Probability of licking on wrong trials, OT, Monkey B (left): P = 0.6208; ranksum test; Monkey S (right): P = 0.9908; t-test j. Probability of licking on wrong trials, learning, Monkey B (left): P = 0.2711; t-test; Monkey S (right): P = 0.8054; paired t-test. k. Visual exploration on wrong trials, OT, Monkey B: P = 0.5932; ranksum test; Monkey S: P = 0.6048; ranksum test l. Visual exploration on wrong trials, learning, Monkey B: P = 0.2424; ranskum test; Monkey S: P = 0.1058; paired t-test.

**Figure S8:**
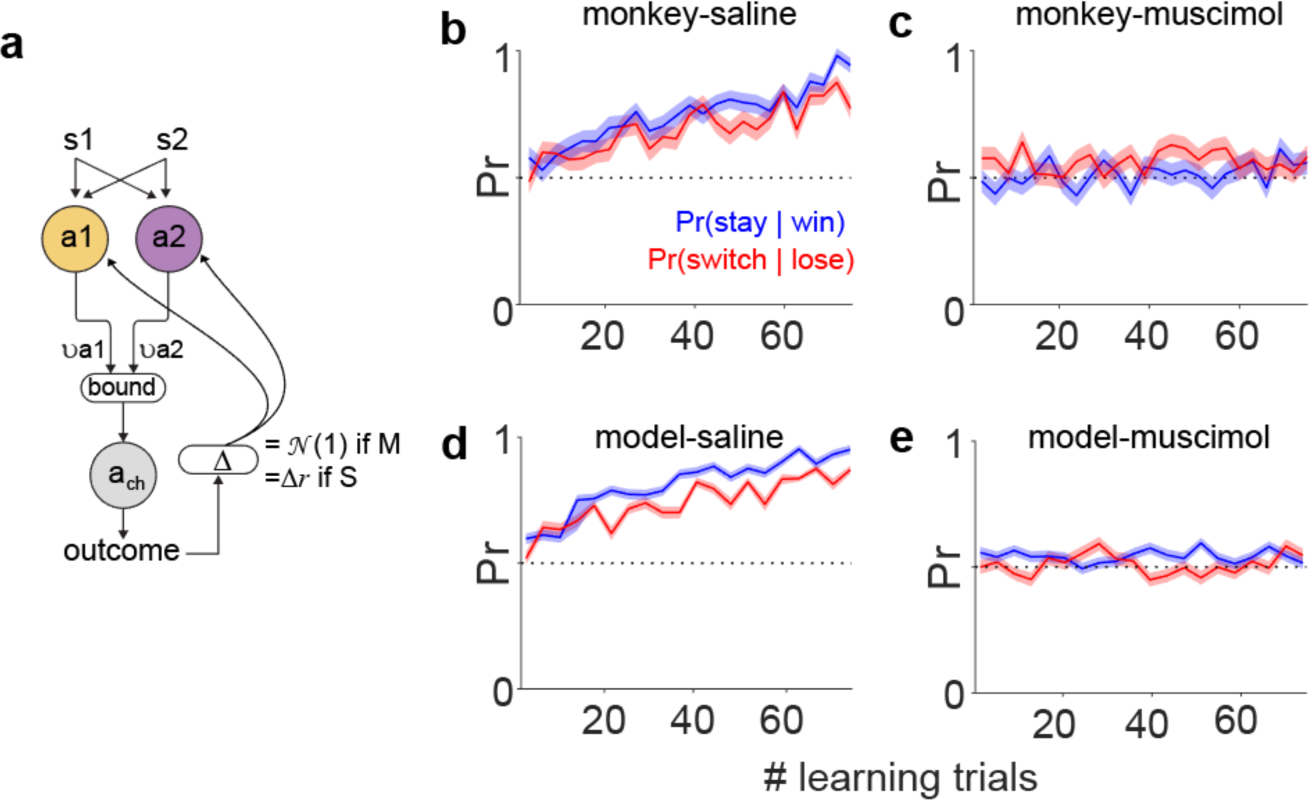
Cerebellar inactivation perturbed the win stay lose switch strategy. a. Reinforcement-drift diffusion model. Action choices are modeled as accumulators with rates ν_a1_ and ν_a2_, racing to threshold (bound). The winner takes all and consequences of the chosen action a_ch_ is evaluated by the activity of P-cells given by Δ. This is used to update the rates of the accumulator on a trial by trial basis. For saline-control condition, Δ is given by the difference in firing rate of correct and wrong outcome in the delta epoch of the P-cells (Δr). For muscimol condition, this is a random number drawn from a Gaussian distribution. b. Probabilities of win-stay and lose-switch during learning from both monkeys during saline-control condition c. Same as **Fig S8a** but during cerebellar inactivation condition. d. Reinforcement learning model prediction for saline-control condition with intact P-cell delta activity. e. Reinforcement learning model prediction for Cerebellar inactivation condition with absent P-cell delta activity.

**Figure S9:**
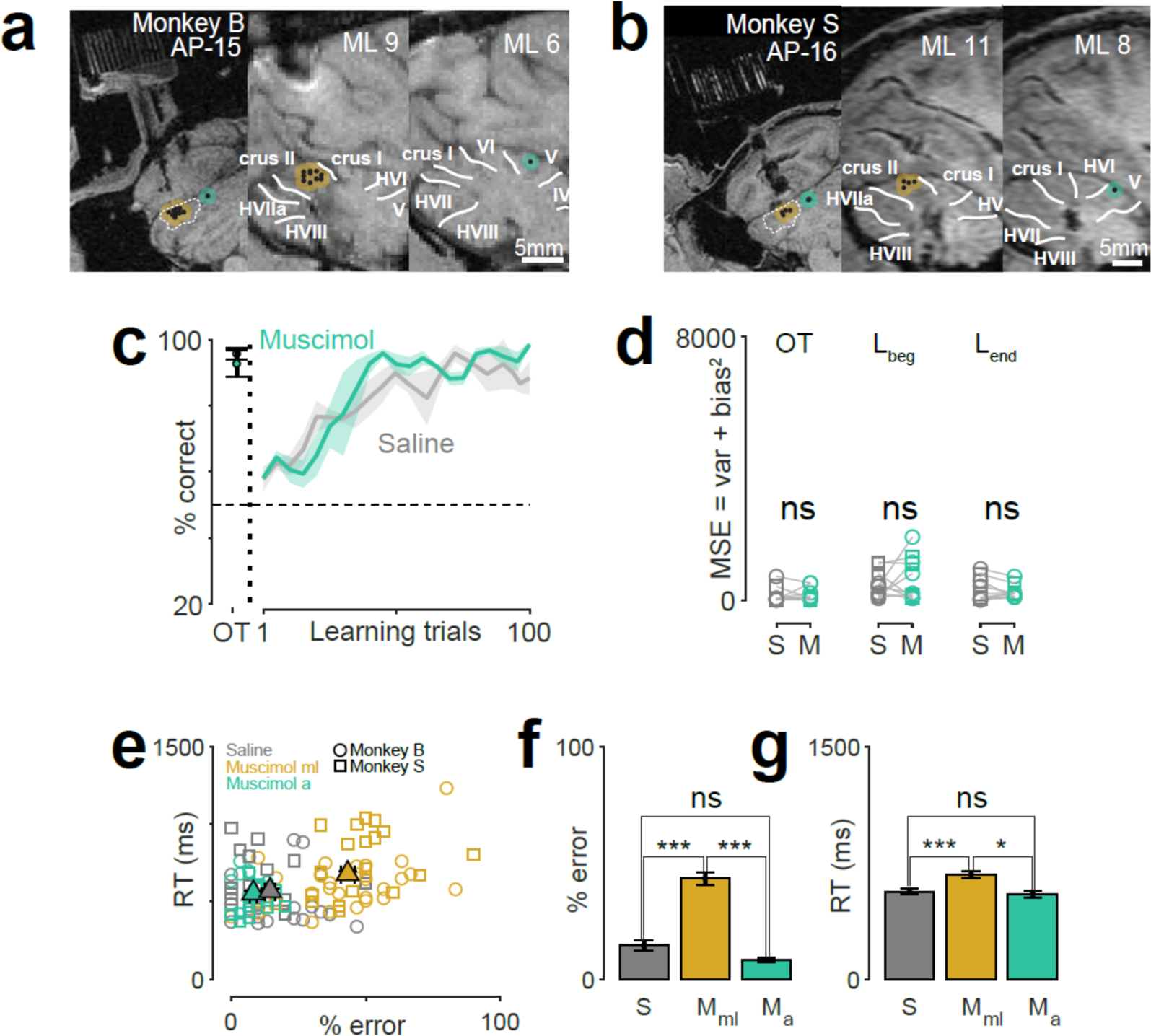
Inactivation of anterior cerebellum did not affect monkey’s performance. a. T1 MRI showing injection regions in the mid-lateral cerebellum (yellow) and anterior cerebellum (green) for monkey B. Same format as **Fig S1**. b. Same as above but for monkey S. c. Behavioral performace of both nonkeys during OT and learning trials for control-saline (gray) and anterior cerebellar inactivation (green) conditions. d. Quantitation of mean squared error (MSE = bias^2^ + variance) from **Fig S9c**; e. RT vs % error for saline injections (S; gray), muscimol injections in the mid-lateral cerebellum (M_ml_; yellow) and anterior cerebellum (M_a_; green). Each maker is an individual session (for one repetition of association learning). Triangles with error bars are means and s.e.m. f. The % error between S and M_ml_ was significant (1.1441e-10; ranksum test) and so was between M_a_ and M_ml_ (2.0153e-09; ranksum test) but not between S and M_a_ (0.1217; ranksum test) g. The RT between S and M_ml_ was significant (1.5543e-04; ranksum test) and so was between M_a_ and M_ml_ (0.0159; ranksum test) but not between S and M_a_ (0.8732; ranksum test)

## Methods

### Animal subjects and surgery

We have described all procedures in detail elsewhere ^4^. We describe them here briefly. We used two male adult rhesus monkeys (*Macaca mulatta*), B and S, for all the experiments. All experimental protocols were approved by the Animal Care and Use Committees at Columbia University and the New York State Psychiatric Institute, and complied with the guidelines established by the Public Health Service Guide for the Care and Use of Laboratory Animals. We located the cerebellum in each monkey using a Simons 3T magnet. We localized the mid-lateral cerebellum using Brainsight, Rouge research Inc. and accessed them through two recording cylinders, on the left hemisphere of monkey B and right hemisphere of monkey S.

### Tasks

We used the NIH REX system for behavioral control^28^. The monkey sat inside a dimly lit recording booth, with its head firmly fixed, in a Crist primate chair 57 mm in front of a LCD monitor upon which visual images were presented which were controlled by a PC running the NIH VEX graphic system.

#### Two-alternative forced-choice discrimination task

The task began with the monkeys grasping two bars, one with each hand, after which a white 1° x 1° square appeared as a trial cue for 800 ms. Then one of a pair of symbols appeared, briefly for 100 ms in some sessions or until the monkey initiated a hand response in other sessions, at the center of gaze. One symbol signaled the monkey to release the left bar and the other to release the right bar. We rewarded the monkeys with a drop of juice for releasing the hand associated with that symbol. The liquid reward and a beep paired with opening of the solenoid were delivered immediately (with a delay of 1 ms) after the initiation of the correct hand movement. We trained the monkeys to associate a specific pair of symbols (green square and pink square) with specific actions (left and right-hand release, respectively) for about 4-6 months until their performance was above 95% correct; we refer to this as the overtrained condition.

In the visuomotor associative learning version of the task (**Fig 1a**), we began every recording session by presenting the monkeys with the same overtrained symbol pair (overtrained condition), and after a number of trials, switched the symbol pair to two fractal symbols, which the monkey had never seen before (novel condition). Both the symbols had equal probabilities of occurring in every trial. The manipulanda remained the same throughout the task. We used two versions of the visuomotor associative learning task shown in **Fig S3**. In the more difficult version, the symbols appeared with equal probability regardless of the monkey’s prior decision. In the easier version, we used an error-correction strategy, showing the same symbol until the monkey made a correct decision. Monkeys learned the symbol-hand association faster during this paradigm, under control conditions. The manipulanda remained the same throughout the task.

In the manipulanda change task (**Fig S6**), we began every recording session by presenting the monkeys with the same over-trained symbol pair and bar manipulanda, and after a number of trials, switched the bar manipulanda to dowel manipulanda. The visuomotor association remained the same

### Data collection and analysis

#### Infusion protocol

At the beginning of each recording session, we lowered a cannula filled with 10 μL of either saline or 10 μg/μL muscimol (diluted in 1X phosphate buffered saline solution), in a track selected on the basis of previous recordings near crus I and II of the mid-lateral cerebellum. We slowly infused the solution using a syringe pump (Harvard Apparatus) through a direct line to the cannula at a constant rate (0.2-0.5 μL/min for 20-30 min). Therefore, we delivered a total mass of 4-10 μg of muscimol in each session. Previous studies have reported that muscimol diffuses and hence functionally inactivates the neurons in an estimated span of 2-3 mm of spherical radius^29,30^. Infusions were typically made at different depths within a single track (total span of ∼ 5 mm) to increase the diffusion radius of the chemical. After infusion, the cannula was left for at least 20-30 min *in situ.* Occasionally, we lowered an electrode in the same track at the same depth after the chemical infusion to confirm silencing of neurons.

#### Single unit recording

We introduced glass-coated tungsten electrodes with an impedance of 0.8-1.2 MOhms (FHC) into the left mid-lateral cerebellum of monkeys every day that we recorded using a Hitachi microdrive. We passed the raw electrode signal through a FHC Neurocraft head stage, and amplifier, and filtered through a Krohn-Hite filter (bandpass: lowpass 300 Hz to highpass 10 kHz Butterworth), then through a Micro 1401 system, CED electronics. We used the NEI REX-VEX system coupled with Spike2 (CED electronics) for event and neural data acquisition. We verified all recordings off-line to ensure that we had isolated Purkinje cells^4^.

#### Analysis of the learning behavior

We constructed the learning curve for every session by calculating the percent correct trials in a sliding window of 10 trials as the bin width moved by 5 trials. Then, we averaged the learning curves across repeated sessions for each symbol pair for each monkey (16 for monkey B and 9 for monkey S) to obtain a total of 25 learning curves. We estimated when the performance first increased above 75% and referred to it as the acquisition-difficulty level for each session. We repeated this for both saline-control condition and the muscimol-inactivation condition.

#### Hand tracking

We painted a spot on the monkeys’ right hand with a UV-blacklight reactive paint (Neon Glow Blacklight Body Paint) prior to every session. We used a 5W DC converted UV black light illuminator to shine light on the spot. Then we used a high speed (250 fps) camera (Edmund Optics), mechanically fixed to the primate chair, to capture a video sequence of the hand movement while the monkeys performed the tasks. We used the track mate feature^31,32^ and custom written software in MATLAB to semi-manually track the fluorescent paint spot painted on the monkey’s hand ^4^.

#### Licking

We recorded licking at a sampling rate of 1000 Hz using a capacitive touch sensor coupled to the metal water spout which delivered liquid water reward near the monkey’s mouth. Raw binary lick traces were used to generate instantaneous lick rate by trial averaging and smoothing it with a Gaussian kernel of sigma = 20 ms.

#### Eye movements

We tracked the monkey’s left eye positions at 240 Hz sampling rate with an infrared pupil tracker (ISCAN, Woburn, MA USA) interfaced with Spike 2 (CED electronics) where it was upsampled to 1000 Hz and synced with the event markers from NEI REX-VEX system. To calculate the radius of visual exploration (r_exp_), we first create a heatplot with overlayed eye movements (**Fig 3e-f**). We then fit a 2D guassian distribution to this data to see the extent of spread from the center and calculated the full-width-at-half-maximum.

#### Reinforcement-drift diffusion model

Details of the model are described elsewhere^4^. We briefly describe it here. Consider learning one symbol-action-outcome association for one symbol with two alternative action choices, *a*_*L*_ and *a*_*R*_ corresponding to left and right hand bar-release hand movements respectively. We model the action selection through a race to threshold model where there is a race in the evidence accumulation between the action values *a*_*L*_ and *a*_*R*_ modeled by Wiener first-passage time (WFPT) distribution. This calculates the likelihood of the reaction time of choice *i* given by:

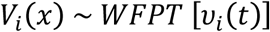

where *v*(*t*) denotes the rate of accumulation process for the trial *t*. The rate of accumulation in the overtrained (OT) condition, 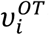 is assumed to be a constant. Then, we model the evolution of *v*_*i*_(*t*), with learning as:

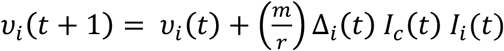

where

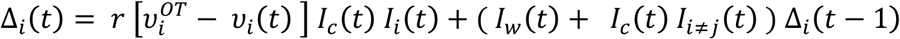

here, *m* captures the rate of learning, *r* is the scale factor, *I*_*i*_(*t*) is the indicator function describing the chosen action (*a*_*ch*_) on trial *t* defined as follows:

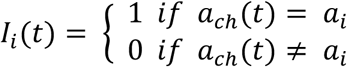

We propose that the rate of change of the accumulation rate *v*_*i*_(*t*), given by Δ_*i*_(*t*) is represented in the delta epoch of the P-cell population that fired at the time of interest. We estimated the accumulator’s initial and final rates by fitting a drift-diffusion model to the observed RT values. Then, we minimized the error between the generated and the empirical RT distributions for the OT and initial learning conditions to get estimates of the parameters. We then simulated the learning process with these values and calculated the probability of win-stay and lose-switch as a function of learning (**Fig S8**). For cerebellar inactivation condition, we modeled the firing rate of P-cells in the delta epoch, Δ_*i*_(*t*) to be a random Gaussian noise and repeated all the above steps otherwise identically (**Fig S8**).

#### Statistics

To check if two independent distributions were significantly different from each other, we first performed a two-sided goodness of fit Lilliefors test, to test for normality, then used an appropriate t-test; or else a non-parametric Wilcoxon ranksum test. All error bars and shading in this study, unless stated otherwise, are mean ± s.e.m.

